# Precise and pervasive phasic bursting in locus coeruleus during maternal behavior

**DOI:** 10.1101/2021.03.31.437751

**Authors:** Roman Dvorkin, Stephen D. Shea

## Abstract

The noradrenergic locus coeruleus (LC) mediates key aspects of arousal, memory, and cognition in structured tasks, but its contribution to natural behavior remains unclear. Neuronal activity in LC is organized into sustained (‘tonic’) firing patterns reflecting global brain states and rapidly fluctuating (‘phasic’) bursts signaling discrete behaviorally significant events. LC’s broad participation in social behavior including maternal behavior is well-established, yet the temporal relationship of its activity to sensory events and behavioral decisions in this context is unknown. Here, we made electrical and optical recordings from LC in female mice during maternal interaction with pups. We find that pup retrieval stably elicits precisely timed and pervasive phasic activation of LC that can’t be attributed to sensory stimuli, motor activity, or reward. Correlation of LC activity with retrieval events shows that phasic events are most closely related to subsequent behavior. We conclude that LC likely drives goal-directed action selection during social behavior with globally-broadcast noradrenaline release.

---

Neurons in the midbrain nucleus locus coeruleus (LC) release noradrenaline (NA) broadly throughout the central nervous system, constituting a key regulator of emotion, arousal, stress, and memory ^1^. Classically, temporal patterns of firing in LC are understood to be composed of two distinct processes: slowly evolving, sustained (‘tonic’) firing, which is thought to reflect global brain states such as arousal, and rapidly fluctuating, bursty (‘phasic’) firing ^2^. Phasic bursts are typically associated with highly salient stimuli and events ^3–5^ or shifts in behavioral strategy and task contingencies ^6–8^. While early work interpreted phasic bursts in LC as responses to behaviorally significant sensory events, over the past two decades, it has become clear that LC neural activity is more precisely aligned with subsequent execution of conditioned behavioral responses ^9–13^. Phasic bursts are therefore perhaps better thought of as linking salient sensory events with learned actions ^2^. The spatial extent and temporal precision of coordinated phasic firing among individual LC neurons remains a subject of investigation ^14–16^, but they may be highly dependent on behavioral context ^17^. Nevertheless, based on their relationship to significant events in structured tasks, LC phasic bursts are an established participant in shaping goal-directed behavior.

Seemingly independent from this central role in navigating experimenter-orchestrated tasks, LC and NA have long been linked to a number of natural social behaviors. For example, microdialysis measurements reveal that NA undergoes elevated and sustained release during conspecific encounters, such as mating ^18^. In the main and accessory olfactory systems, NA is essential for establishing memories of mating partners ^18–21^, for maternal bonding and subsequent recognition of offspring in sheep ^22^, and for imprinting offspring to odors associated with maternal care ^23^. Female mice lacking the enzyme dopamine-beta-hydroxylase (Dbh), essential for synthesizing NA from dopamine, showed profound disruption of maternal behaviors, often resulting in pup death due to neglect ^24^. Importantly, in that study, maternal care was restored in *Dbh^-/-^* mutant mothers when NA synthesis was reactivated shortly before birth.

Collectively, these observations compellingly inculpate NA and LC in a range of natural social behaviors, including maternal care and motivation in particular. However, the technical approaches used lacked sufficient spatial and temporal resolution to ascertain the timing and structure of the underlying neuronal activity in LC. Therefore, the relative contributions of tonic and phasic firing patterns to social interactions are unknown, and the timing of these patterns relative to salient social events is undetermined. Specifically, it is unclear whether fluctuations in NA release occur in response to specific social sensory stimuli, or whether they anticipate specific goal-directed social behaviors.

Here we used chronic *in vivo* electrophysiology and fiber photometry to measure single unit and population neural activity in LC of freely behaving surrogate mice during their interactions with pups. The term ‘surrogate’ here refers to a virgin female who is co-housed with a mother and her pups beginning before birth, and who learns to perform maternal care over the first few postnatal days. We found that several aspects of maternal care were reliably associated with fluctuations in LC activity. Contact with pups during retrieval events precisely coincided with phasic bursts in individual LC neurons and rapid, transient increases in optically detected bulk fluorescence that continued until the pup was dropped in the nest. The ubiquity of this response among LC neurons, and its reliability and magnitude in fiber photometry recordings, strongly suggest that these events are coordinated across LC and broadcast NA release throughout the brain. We also observed slow changes in tonic firing rate when females performed distinct maternal behaviors such as nest building and pup grooming. Retrieval-related LC bursts could not be explained merely by responses to sensory stimuli, general motor activity, or reward, and changes in tonic firing were not seen during highly similar, but non-social motor activities. Analysis of the relationship between phasic events and retrieval behavior indicates that LC activity specifically correlates with impending behavior. We conclude that rapid changes in LC firing likely regulate social behavior in part by promoting specific context-dependent actions.

## RESULTS

### Individual LC neurons emit brief phasic bursts locked to pup retrieval

To directly observe the firing of noradrenergic LC neurons (LC-NA) during maternal interaction, we chronically implanted miniature movable drives carrying a bundle of 8 or 16 microwires targeted to a location 1 – 0.5 mm above LC in nulliparous female mice (n = 3). Over a period of approximately two weeks, drives were lowered daily by 50 – 100µm until putative LC-NA were located based on well-established electrophysiological criteria (see Methods) ^1, 21, 25^. During this time, mice were habituated to the experimental arena and to the experimenter with gentle handling. Immediately following the initial identification of putative LC-NA, each subject was co-housed with a female in late-stage pregnancy. After several days in these housing conditions, nulliparous females begin to exhibit characteristic maternal behaviors, including retrieval of pups who become separated from the nest and emit ultrasonic distress vocalizations (USVs) ^26–30^. Following birth, the now maternally-experienced females (‘surrogates’) were placed with the familiar pups in the experimental arena for extended sessions of free interaction (40 – 90 min). During each session, neural activity was detected and recorded by a lightweight flexible cable tethering the animal to a motorized commutator. Neural spiking signals were manually sorted offline (OpenEx, TDT) and every recording site reported here met criteria to be designated as a single unit (see Methods).

Pup retrieval was elicited several times during each session when the experimenter scattered the pups across the arena and waited for the surrogate to return them to the nest (Fig. 1A). By doing this, we were able to collect LC neural data associated with numerous individual retrieval events from each session (mean: 21.2; range: 5 – 30). Twelve putative LC-NA reliably exhibited phasic bursts close to the time of each retrieval event (Fig. 1B). These bursts strongly resembled the typical burst-pause response seen in LC-NA in response to significant or surprising events such as a tail pinch ^25^. To determine the timing of these bursts for each neuron relative to each retrieval, we compared the alignment of the spike train for each trial from a neuron to three different components of retrieval (pup contact, pup lift, pup drop Fig. 1C-E; Extended Data Fig. 1). Once each neuron was aligned, we compared the PSTH width at half-maximum (Fig. 1F) and measured the mean lead or lag of the neuronal activity relative to the specific behavior (Fig. 1G). When aligned to contact, the half-max width of the PSTH was 1.79 ± 0.22 s and the peak of firing lagged by 1.10 ± 0.23 s. (Unless otherwise stated, all values are mean ± SEM). When aligned to lift, the half-max width of the PSTH was 1.98 ± 0.22 s and the peak of firing lagged by 0.56 ± 0.18 s. When aligned to drop, the half-max width of the PSTH was 2.51 ± 0.54 s and the peak of firing led by 0.98 ± 0.27 s. These values for pup drop were significantly different from those for contact and lift (paired t-test; *p* < 0.05). Based on the narrow peak seen in the PSTH, and the fact that the firing was centered around the mouse lifting the pup, for the remainder of this study, we will use the term ‘retrieval’ to refer to this point. When we aligned to this point, we found that individual bursts were narrow but exhibited different phases relative to retrieval; some neurons increased firing immediately following contact and others at later times during the mother’s return trip to the nest. All neurons but one exhibited a characteristic inhibition (pause) just after pup drop (Fig. 2A, Extended data Fig. 1).

**Figure 1:**
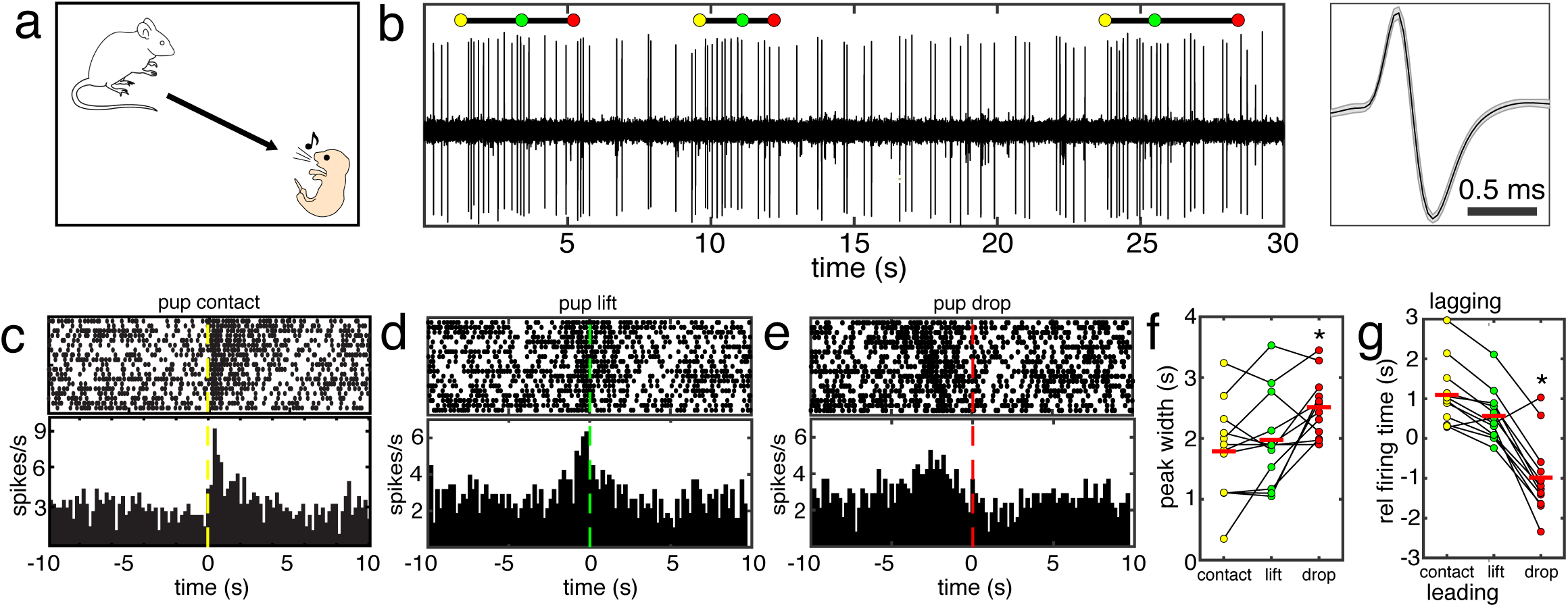
Noradrenergic LC neurons (LC-NA) emit brief phasic bursts timed to pup retrieval. (a): Schematic of the behavior. The pup emits USVs, summoning the female to pick it up and return it to the nest. (b): Example single unit data. *Left panel:* A trace of 30 s of a recording from an LC-NA neuron that includes three retrieval events. Three key components in the behavioral sequence are marked with colored dots: yellow = pup contact, green = lifting the pup (retrieval onset), and red = the time the pup was dropped in the nest. *Right panel:* The mean of 100 spikes from the neuron depicted on the left. The gray shaded band around the mean is the SEM. (c): Raster plot and peristimulus time histogram (PSTH) of neural data recorded from the neuron in (b) during 19 retrieval events from one session. Data in these plots are aligned to the contact of the female with the pup. (d): Same as (c), but the data are aligned to time that the female lifts the pup. (e): Same as (c), but the data are aligned to the time that the female drops the pup in the nest. (f): Scatterplot comparing the width of the peak in the PSTH at half-max height above baseline between data aligned to contact, lift, and retrieval. (contact: 1.79 ± 0.22 s; lift: 1.98 ± 0.22 s; drop: 2.51 ± 0.54 s). The width of the peak was significantly greater when spikes were aligned to drop as compared to contact or retrieval (paired t-test; *p* < 0.05). (g): Scatterplot comparing the lag (+) or lead (−) of the peak in the PSTH relative to contact, lift, and drop. (contact: 1.10 ± 0.23 s; lift: 0.56 ± 0.18 s; drop: −0.98 ± 0.27 s). The lag of the peak was significantly different for drop as compared to contact or lift (paired t-test; *p* < 0.05).

**Figure 2:**
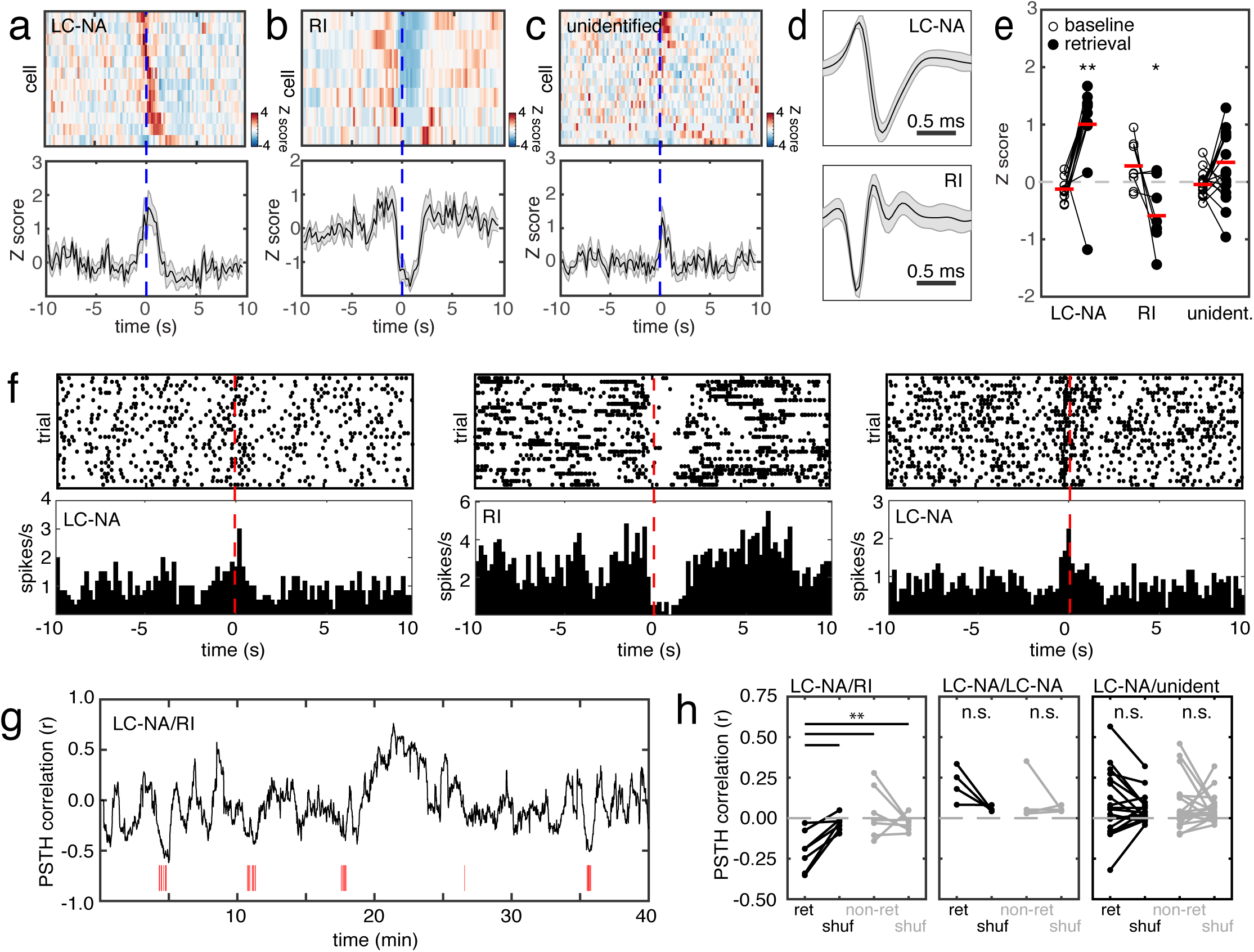
Physiologically distinct local neurons show activity that is inversely correlated to LC-NA during retrieval. (a): A 2D PSTH of the mean activity of all LC-NA (n = 12) during retrieval (*top panel*) and a plot of the mean Z-score firing rate over all cells (*bottom panel*). In the top panel, each row is the mean firing rate for one neuron, and the data are aligned to lift (retrieval). (b): Data from all retrieval inhibited (RI) neurons (n = 7) organized as in (a). (c): Data from all unidentified neurons (n = 18) organized as in (a). (d): Examples of typical spike shapes from LC-NA (*top panel*) and RI (bottom panel). Each plot is the mean of 100 spikes from the corresponding neuron. The gray shaded band around the mean is the SEM. (e): Scatterplot comparing the mean activity during retrieval to the immediately preceding baseline activity for all neurons of all three types. LC-NA showed significantly higher firing rates during retrieval than at baseline (n = 12; baseline: −0.13 ± 0.06 Z-score; retrieval: 1.00 ± 0.23 Z-score; paired t-test, *p* < 0.01). RI showed significantly lower firing rates during retrieval than at baseline (n = 7; baseline: 0.28 ± 0.15 Z score; retrieval: −0.59 ± 0.20 Z-score; paired t-test, *p* < 0.05). (f): Raster plots and PSTHs of neural data recorded simultaneously from three neurons (two LC-NA and one RI) during 28 retrieval events from one session. Data are aligned to lift (retrieval). (g): A plot of the fluctuating correlation (see Methods) of one simultaneously recorded LC-NA/RI pair. The red tick marks denote the times of retrieval. The strongest negative correlations were seen during clusters of retrieval events. (h): Scatterplots comparing the mean correlation values for all three neuron types paired with simultaneously recorded LC-NA. Correlation values are plotted separately for firing during retrieval episodes and non-retrieval activity, and all values are also computed for shuffled data. Correlation values were significantly more negative in LC-NA/RI pairs during retrieval than in non-retrieval periods or shuffled data (n = 7; retrieval: −0.21 ± 0.5; non-retrieval: 0.00 ± 0.05; shuffled retrieval: −0.05 ± 0.02; one way ANOVA, *p* < 0.01).

### Faster spiking neurons in or near LC are sharply inhibited during pup retrieval

In the course of performing these experiments, we incidentally recorded other single units in the immediate vicinity of LC which did not match the characteristics of LC-NA (n = 25). For example, post hoc analysis revealed a group of bursty, faster spiking (8.55 ± 1.2 spikes/s) neurons with a narrow shape (Fig. 2D) which were consistently and strongly inhibited around the time of pup retrievals (retrieval inhibited, ‘RI’ neurons). These neurons consequently exhibited a reciprocal response to LC-NA including a strong rebound in firing rate just after the time the pup was dropped in the nest (n=7 neurons from 2 mice; Fig. 2C, D; Extended Data Fig. 1). Alignment to contact revealed a sharp increase in the firing rate of these neurons beginning a few seconds prior to and peaking at contact (Extended Data Fig. 1B). Closer examination of our retrieval videos revealed that this corresponded to the mouse initiating motion towards the pups.

In addition, we collected another 18 neurons that did not match the profiles of LC-NA or RI; these neurons were designated ‘unidentified’ and rarely exhibited activity related to retrieval (Fig. 2C, Extended Data Fig. 1). We quantified the sign and magnitude of retrieval activity in each of these three classes by converting the firing rates of all neurons to Z-scores and comparing the mean baseline activity just before retrieval to the mean activity during retrieval (Fig. 2E). Putative LC-NA showed a significant increase in firing rate during retrieval (n = 12; baseline: −0.13 ± 0.06 Z score; retrieval: 1.00 ± 0.23 Z-score; paired t-test, *p* < 0.01), while RI neurons showed a significant decrease in firing rate during retrieval (n = 7; baseline: 0.28 ± 0.15 Z score; retrieval: - 0.59 ± 0.20 Z-score; paired t-test, *p* < 0.05). As a group, the unidentified neurons did not show a significant change in firing (n = 18; baseline: −0.05 ± 0.05 Z score; retrieval: ± 0.23 Z-score; paired t-test, p = 0.028 did not survive the correction for multiple comparison).

We were not able to definitively identify the location of the cell bodies of the RI neurons, however, the apparently reciprocal firing pattern during pup retrieval raises the possibility that RI correspond to GABAergic neurons in and around LC that have been proposed to exert inhibitory control of LC-NA ^31–34^. Many of our recording sessions yielded multiple simultaneously recorded neurons (Fig. 2E). We therefore examined correlated activity between LC-NA and all three types of neurons (other LC-NA, RI, and unidentified; 30 neuron pairs) (Fig 2F, G). We binned firing rate for both neurons in each pair at 1 s/bin and then performed a Pearson correlation on the bin values of the two histograms.

Correlations between LC-NA and RI fluctuated in time, typically reaching maximum inverse correlation during episodes of retrieval (Fig. 2F). We therefore separately computed the correlations for bins falling in the 8 s following a retrieval event from those falling elsewhere in the record (Fig. 2G). Mean correlation value for LC-NA/RI pairs during retrieval episodes was significantly more negative than that for all other time points (n = 7 pairs; retrieval: −0.21 ± 0.05; non-retrieval: 0.00 ± 0.05; paired t-test, *p* < 0.05). The inverse correlation of these two cell types during retrieval was not due to a non-specific synchronizing effect of retrieval because it disappeared when retrieval trials were shuffled between the paired neurons (n = 7 pairs; retrieval: −0.21 ± 0.05; shuffled retrieval: −0.05 ± 0.02; paired t-test, *p* < 0.01). Mean correlation value for LC-NA/RI pairs during non-retrieval periods did not significantly differ from that obtained when the temporal shift between the spike trains of the paired neurons was circularly permuted (n = 7 pairs; non-retrieval: 0.00 ± 0.05; shuffled non-retrieval: 0.00 ± 0.0; paired t-test, *p* = 0.95).

No significant differences in correlation values were seen between retrieval and non-retrieval data, or between shuffled and non-shuffled data, for LC-NA/LC-NA pairs or for LC-NA/unidentified pairs.

### Optical recordings reveal that retrieval activity in LC is pervasive and synchronous

Our electrophysiology data have two limitations. First, mounting evidence argues that LC may have a more modular organization anatomically and functionally than was previously appreciated ^15, 16, 35, 36^. Hence, despite the fact that all of our putative LC-NA exhibited sharply elevated phasic activity during pup retrievals, it is possible that LC neurons projecting to different parts of the brain fire differently during pup interaction. Second, due to the challenging nature of electrical recordings in freely behaving mice, data are sporadic and low yield, making longitudinal experiments nearly impossible. To overcome these limitations, we used fiber photometry to measure bulk Ca^2+^ signals from LC-NA. This method captures the neuronal population activity, so if the timing of LC-NA participation in pup retrieval is heterogenous, the signal fluctuations will presumably be of relatively low amplitude. Fiber photometry also allows us to monitor the same neural population day by day, so we can monitor any changes in the responses as the surrogate learns to perform maternal behavior.

We injected a Cre-dependent AAV carrying the genetically-encoded Ca^2+^ sensor GCaMP6s into the LC of *Dbh-Cre* mice, thereby restricting GCaMP expression to noradrenergic neurons (Fig. 3A-C). After 3-5 weeks to allow for full viral expression and recovery, surrogates were co-housed with a pregnant female. Following birth, retrieval experiments were conducted over several consecutive days in a familiar experimental arena. Consistent with what we observed in single unit recordings from putative LC-NA, pup retrievals were accompanied by a steep, strong rise in the population Ca^2+^ signal (Fig. 3D) beginning around contact with the pup and reaching its peak just after retrieval initiation (Fig. 3F) (n = 7 mice; baseline: 0.025 ± 0.07 Z-score; retrieval: 1.36 ± 0.13 Z-score; paired t-test, *p* < 0.001). The signal remained elevated through the retrieval until the pup was dropped in the nest. Signals typically subsequently exhibited a dip below baseline (n = 7 mice; post drop dip: −0.50 ± 0.13 Z-score; paired t-test compared to baseline, *p* < 0.05) that resembled the burst-pause response typical of phasic activity in LC-NA ^25^ (Fig 3G, H). Comparing the signal across 5 consecutive days, we found no significant changes in the relative amplitudes and timing of the Ca^2+^ peaks (Fig. 3I, J) (one-way ANOVA, *p* > 0.05).

**Figure 3:**
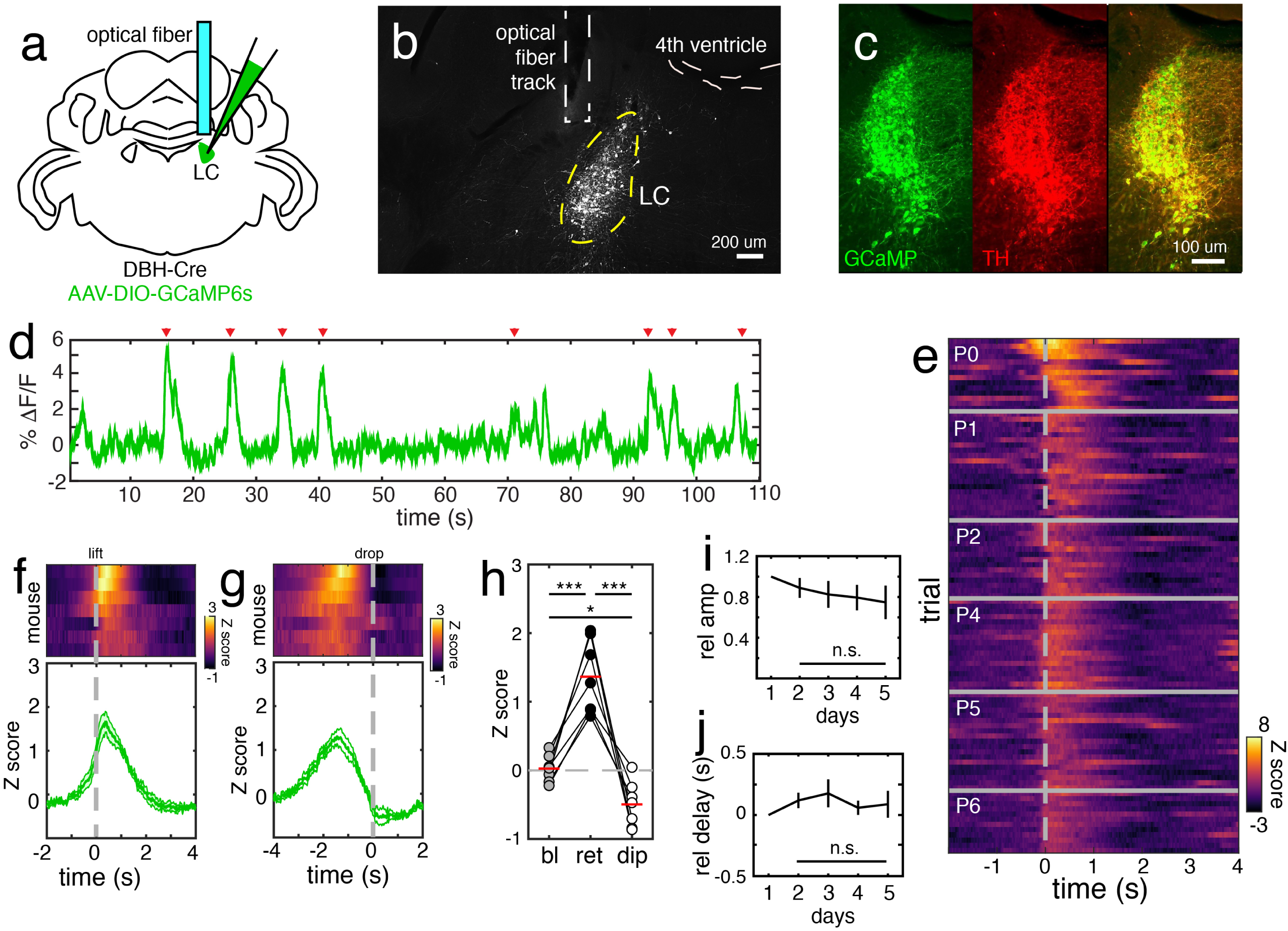
Fiber photometry reveals that LC-NA emit pervasive and synchronous population bursts during retrieval. (a): Experimental strategy. DBH-Cre mice were injected with a Cre-dependent AAV driving expression of the genetically-encoded Ca^2+^ indicator GCaMP6s and were implanted ipsilaterally with an optical fiber. (b): Photomicrograph showing GCaMP expression and the location of the fiber. Scale bar = 200 μm. (c): Photomicrographs showing the overlap in expression of GCaMP (green) and tyrosine hydroxylase (TH) (red), a marker of noradrenergic neurons. Scale bar = 100 μm. (d): Plot of 110 s of raw ΔF/F data from one example subject performing 8 retrievals (red arrowheads). Each retrieval is accompanied by a large transient in fluorescence. (e): Heatmap of 99 individual retrieval trials taken from PND0 through PND6 (f, g): Plots of mean responses during pup retrieval for each mouse (n = 7; *top panel*) and the mean ± SEM across all mice, aligned to retrieval (f) and drop (g). (h): Scatterplot comparing the mean Z-score values from baseline, retrieval, and the post retrieval dip for all mice. All values were significantly different from the other values (n = 7; baseline: 0.025 ± 0.07 Z-score; retrieval: 1.36 ± 0.13 Z-score; post drop dip: - 0.50 ± 0.13 Z-score; paired t-test with Holm-Bonferroni correction ***p < 0.001; * p < 0.05). (i, j): Plots of the stability of signals over 5 days for all mice. Mean amplitude relative to day 1 (normalized to 1) (i) and mean timing of peak activity relative to day 1 (defined as 0) are shown for all mice (n = 7).

Pregnancy is associated with hormonal fluctuations that trigger comprehensive changes in the mother’s physiology, including neural circuitry and brain function. Although surrogates learn to perform pup retrieval and other maternal behaviors, it is possible that pregnancy lead to a different LC-NA activation profile as compared to surrogates. We tested this by mating former surrogates expressing GCaMP (n = 3; 1 female was injected with GCaMP6s, 2 females were injected with GCaMP7f) and performing retrievals with their pups beginning after delivery (PND0 – PND5) (Extended Fig. 2). Pup retrieval in mothers elicited strong phasic activation of LC beginning around pup contact and persisting until pup drop (baseline: −0.18 ± 0.04 Z-score; retrieval: 1.03 ± 0.06 Z-score; paired t-test, *p* < 0.001). This activity pattern was qualitatively indistinguishable from the corresponding events in surrogates. Thus, we conclude that LC activity in surrogates is representative of that in mothers.

The strength and precision of the phasic activity in the LC-NA population during pup retrieval leads us to conclude that these events are likely pervasive and reflect synchronous activity among a large proportion of the population of LC-NA. If that is correct, then this specific behavior may evoke coordinated ‘broadcast’ release of NA at many downstream targets.

### LC activation at pup contact predicts subsequent retrieval and is not experience-dependent

We next asked how phasic LC activity during retrieval emerges as the female acquires maternal experience. Naïve virgin female *Dbh-Cre* mice were injected in LC with a Cre-dependent AAV carrying the Ca^2+^ sensor GCaMP7f. Following recovery, the mice were exposed to pups daily (postnatal days 1 – 5) for 30 – 45 min in a familiar experimental arena. Similar protocols have been used to induce maternal behavior in naive rats and mice without the need for continuous co-housing ^37, 38^. At the end of each exposure, pups were scattered across the arena several times as above to elicit maternal retrieval. Initially (PND1 – 2), inexperienced females only investigated pups (transiently touching them with their snout) without initiating retrieval to the nest. These brief contacts did not elicit any detectable increase in population Ca^2+^ signals in LC (Fig. 4A, D) (n = 3 mice; baseline: 0.02 ± 0.1 Z-score; investigation only: 0.03 ± 0.6 Z-score; paired t-test, *p* = 0.90). By PND3, all mice (n = 3) successfully learned to reliably retrieve the scattered pups. Investigatory contact with pups, only when followed by retrieval, was accompanied by a rise in fluorescence that persisted until the pup was dropped in the nest (Fig. 4B) (n = 3 mice; baseline: −0.04 ± 0.03 Z-score; investigation with retrieval: 1.73 ± 0.18 Z-score; paired t-test, *p* < 0.05). Therefore, LC activation in surrogates at the time of initial contact with a pup predicted subsequent retrieval to the nest. Indeed, the activity that ensued following contact appeared more sharply aligned to subsequent retrieval (Fig. 4C) (n = 3 mice; contact alignment: 2.32 ± 0.19 Z-score; retrieval alignment: 2.84 ± 0.15 Z-score; paired t-test, *p* < 0.05). This set of observations is consistent with evidence from structured tasks that phasic LC activity is not driven by sensory stimulation by itself. Rather, bursts in LC are elicited in a context-dependent manner, reliably preceding goal-directed actions made in response to sensory cues ^9–13^.

**Figure 4:**
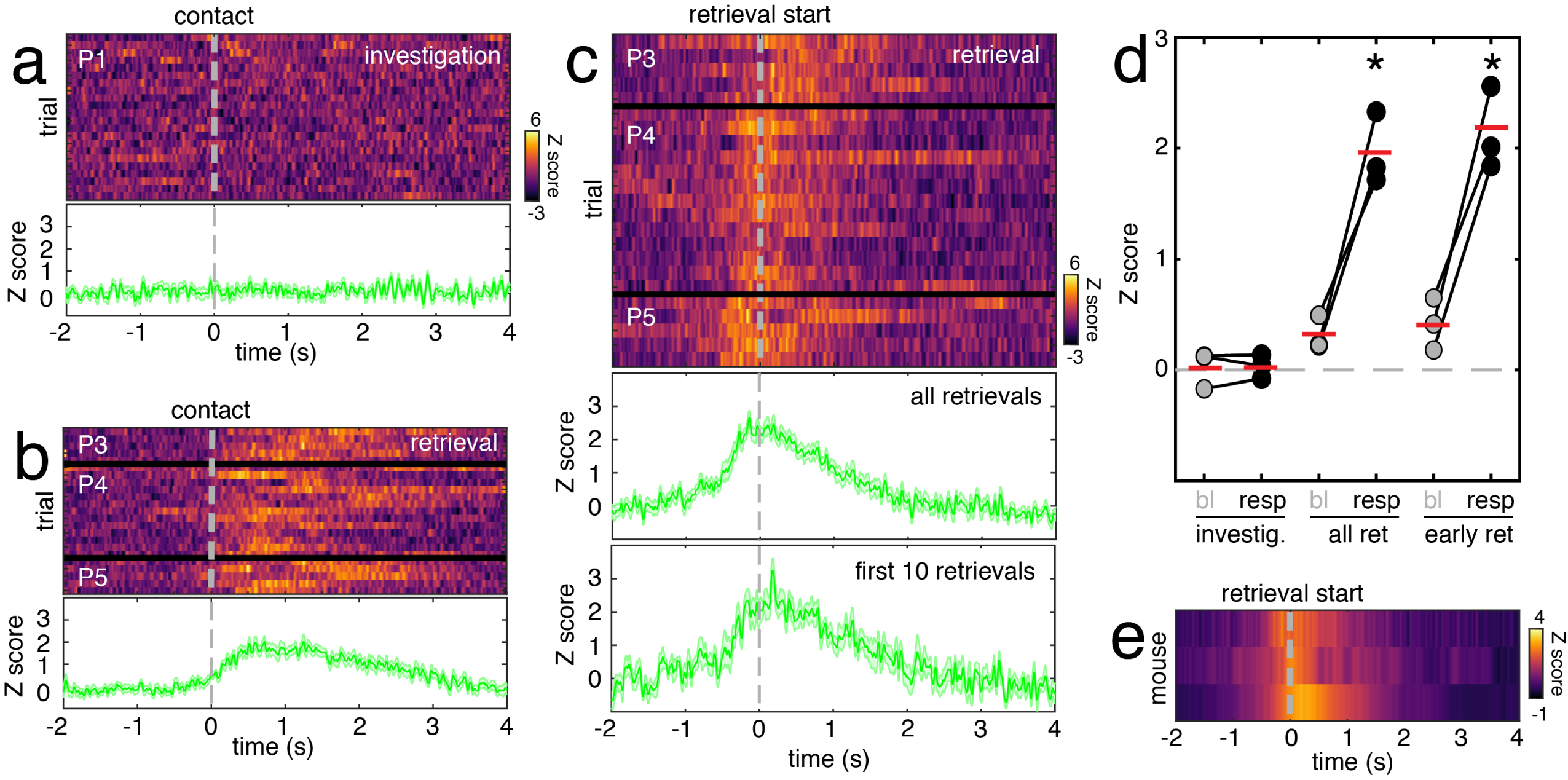
LC activation on contact with pups predicts subsequent retrieval and is not experience-dependent. (a): Heatmap and mean trace showing an example of the lack of response to pup contact when not followed by retrieval. Data were taken from PND1, prior to the emergence of retrieval behavior in a mouse with controlled exposure to pups. We refer to this as investigation. (b): Heatmap and mean trace showing an example of the response to pup contact when it was followed by retrieval. Data were taken from PND3 – 5, after the emergence of retrieval behavior in a mouse with controlled exposure to pups. (c): Heatmap and mean traces showing an example of the response to retrieval as it emerged in a mouse with controlled exposure to pups. Data are aligned to retrieval. All retrievals performed by this mouse are represented in this panel. The mean traces below compare the mean of data from the first 10 retrieval trials exhibited by this mouse to the mean of all trials in the experiment. (d): Scatterplot comparing the baseline activity to the behavior activity for investigation, all retrievals, and the 10 earliest retrievals. Responses during investigation were not significantly different from baseline (n = 3 mice; paired t test; *p* = 0.90). Significant responses relative to baseline were observed for all retrievals as well as for early retrievals only (n = 3 mice; paired t test, *p* < 0.05). The amplitude of early retrievals did not significantly differ from the mean amplitude of all retrievals (n = 3 mice; paired t-test, *p* = 0.17). (e): Heatmap depicting the time course and amplitude of mean retrieval responses for all subjects.

We speculated that the magnitude of phasic activity associated with pup retrieval may gradually increase with improving retrieval performance on early trials, but this was not the case. We compared the mean amplitude of Ca^2+^ signals on the first 10 retrievals performed by each mouse with the mean amplitude of all retrievals from that mouse (Fig. 4C, D). The magnitude of early trials did not significantly differ from all trials (n = 3 mice; early trials: 2.14 ± 0.22 Z-score; all trials: 1.96 ± 0.19 Z-score; paired t-test, *p* = 0.17). This result shows that retrieval-related activation of LC emerges at full amplitude on the very first retrievals performed by the mouse, and is not experience-dependent.

### Pup retrieval activity in LC is distinct from non-social motor or reward responses

The above results seem to indicate that sensory stimuli from the pups (such as pup odors) alone are not sufficient to account for LC activity during pup retrieval. However, it remains possible that the responses are evoked by other, non-social motor or reward aspects of the task. For example, one recent study suggested that LC-NA mediates effort/reward trade-off ^39^. To compare LC responses to non-social motor, effort exerting, and rewarding activities, we measured LC activation patterns during digging, appetitive reward (snack), and retrieval of a toy (fake mouse) using both fiber photometry and single unit electrophysiology.

Lab mice tend to dig in their corncob bedding, looking for pieces to gnaw on. While not directly comparable to pup retrievals, this activity both expends effort and is potentially rewarding. Nevertheless, when we measured LC activity aligned to the onset of digging, there were no significant changes in either Ca^2+^ signal (Fig. 5A, C; baseline: −0.15 ± 0.05 Z-score; digging: −0.09 ± 0.05 Z-score; paired t-test, p = 0.32) or in the firing rate of single units (Fig. 6A, D; baseline: 1.84 ± 1.4 spikes/s; digging: 1.60 ± 1.5 spikes/s; paired t-test, p = 0.30).

**Figure 5:**
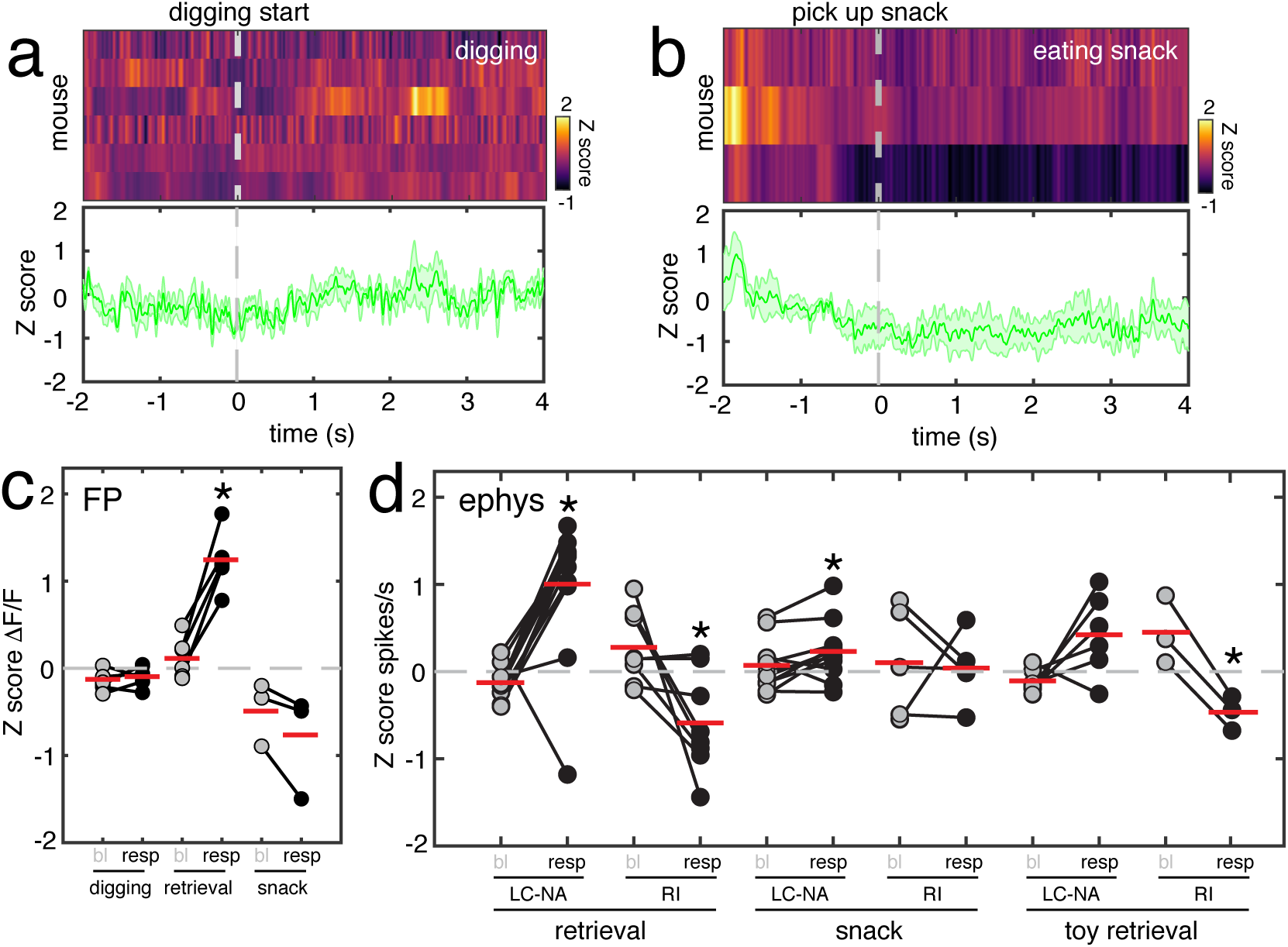
LC maternal retrieval responses are not replicated by motor activity, unexpected reward, or retrieval of an inanimate object. (a): Plots of the mean fluorescence signal in LC aligned to bouts of vigorous digging in the cage bedding. The mean response of LC to the onset of digging in six mice is shown in the heat map (*top panel*). Each row is the mean response of one mouse. The lower plot shows the mean response to digging across all mice. (b): Plots of the mean fluorescence signal in LC aligned to locating a hidden snack buried in the bedding. The mean response of LC to the snack in three mice is shown in the heat map (top panel). The lower plot shows the mean response to snack discovery across all mice. (c): Scatterplot comparing the magnitude of GCaMP fluorescence responses to digging and snack discovery with responses to pup retrieval. In contrast with the robust retrieval responses, neither digging (n = 6 mice; paired t test, *p* = 0.33) nor snack discovery (n = 3 mice; paired t test, *p* = 0.14) resulted in a significant increase above baseline activity. (d): Scatterplot comparing the magnitude of neuronal spiking responses to snack discovery and retrieval of a stuffed mouse toy with responses to pup retrieval for LC-NA and RI neurons. In contrast with the reliable and robust responses in both neuron types to pup retrieval, firing rate changes to the other events were weak and inconsistent. Snack discovery did evoke a significant increase in firing above baseline in LC-NA, but it was far smaller (n = 11 neurons; baseline: 0.07 ± 0.13 Z-score; snack: 0.23 ± 0.15 Z-score; paired t-test, *p* < 0.05). Retrieval of the toy mouse resulted in a significant drop in firing rate only in RI (n = 3 neurons; baseline: 0.45 ± 0.19 Z-score; toy retrieval: −0.47 ± 0.22 Z-score; paired t-test, *p* < 0.05).

**Figure 6:**
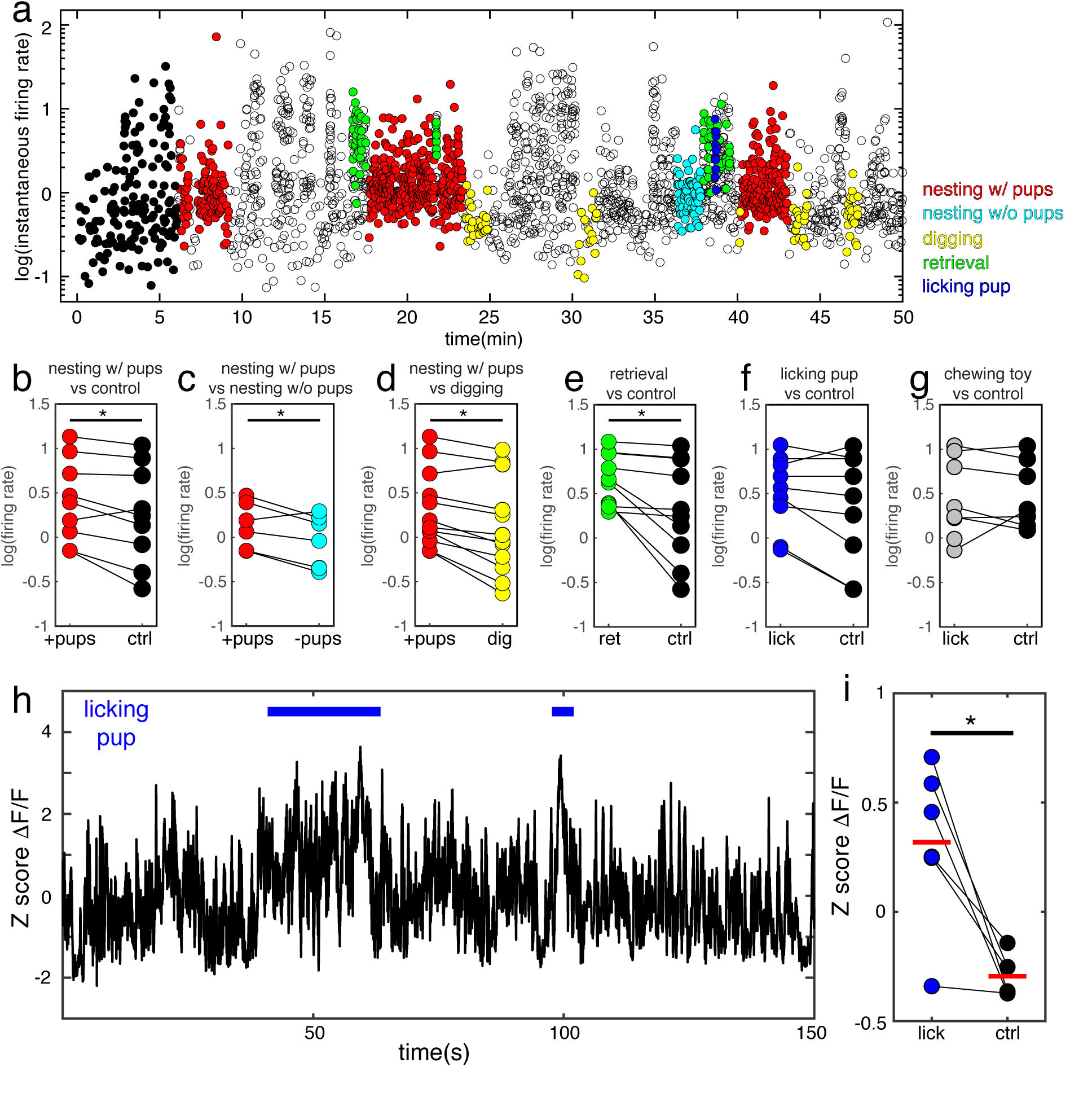
Distinct maternal behaviors are associated with tonic increases in LC neural activity. (a): Plot of log_10_(instantaneous firing rate) from one LC-NA neuron over 50 minutes of recording in a freely behaving pup-experienced female. Some of the points are color-coded to reflect that those data were collected during the specific behavior indicated by the color legend. Nesting with pups refers to active nest building while one or more pups were in the nest. Nesting without pups refers to the same behavior performed without any pups in the nest. The black points found at the start of the trace while the subject was alone in the arena were used as control data to compare with other behaviors. (b-g): Comparison of log firing rates associated with various behaviors, including: nesting with pup vs. control (b), nesting with pups vs. nesting without pups (c), nesting with pups vs. digging (d), retrieval vs. control (e), licking pup vs. control (f), and chewing the toy mouse vs. control (g). Nesting with pups had significantly higher firing rates then control (nesting with pups: 2.53 ± 1.4 spikes/s; control: 1.79 ± 1.5 spikes/s; paired t-test, *p* < 0.05), nesting without pups (nesting with pups: 1.37 ± 1.3 spikes/s; nesting without pups: 0.96 ± 1.3 spikes/s; paired t-test, *p* < 0.05), and digging (nesting with pups: 2.06 ± 1.3 spikes/s; digging: 1.31 ± 1.4 spikes/s; paired t-test, *p* < 0.01). Firing during retrieval was also higher than during control (retrieval: 4.46 ± 1.2 spikes/s; control: 2.07 ± 1.5 spikes/s; paired t-test, *p* < 0.05). (h): Plot of 150 s of GCaMP fluorescence data containing two examples of episodes of the mouse licking and grooming pups, denoted by the blue bars. (i): Comparison of fluorescence signals during licking and grooming of pup and control periods alone in the arena. Activity was significantly higher during licking and grooming (n = 6 mice; licking: 0.32 ± 0.15 Z-score; control: −0.29 ± 0.04 Z-score; paired test, *p* < 0.05).

Introducing a toy mouse to the arena elicited subsequent investigation and, in few cases, retrieval of the toy to the nest. Retrieval of the toy was accompanied by a slight, but not statistically significant increase in firing rate of LC-NA single units as compared to baseline (Fig. 5D; baseline: −0.11 ± 0.05 Z-score; toy retrieval: 0.42 ± 0.19 Z-score; paired t-test, p = 0.08) and a significant decrease in the firing rate of RI units (Fig. 5D; baseline: 0.45 ± 0.19 Z-score; toy retrieval: −0.47 ± 0.22 Z-score; paired t-test, p < 0.05). This partial response could reflect effort associated with retrieval of the toy, however it seems unlikely that the steeper rise in LC-NA firing during pup retrieval is entirely related to predicted effort as the toy was substantially larger than a pup and demanded no less effort to retrieve (Fig. 5D). Moreover, retrieval responses did not increase as pups gained in weight (Figs. 3 and 4).

To test for non-social reward response, mice were given a snack (chocolate pellet). As recently reported ^40^, snack consumption by satiated mice (food and water was provided *ad libitum* to all experimental animals, see Methods) was accompanied by a drop in Ca^2+^ signal that was not significant (Fig. 5B, C; baseline: −0.48 ± 0.21 Z-score; snacks: −0.81 ± 0.35 Z-score; paired t-test, *p* = 0.14). There was a mixed response in the two single unit types (LC-NA and RI) with a minor statistically significant increase in the firing rate of LC-NA units while mice ate the snack as compared to baseline (Fig. 5D; baseline: 0.07 ± 0.13 Z-score; snack: 0.23 ± 0.15 Z-score; paired t-test, *p* < 0.05). These results demonstrate that strong phasic activation of LC during pup retrieval cannot simply be attributed to motor response, effort exertion, or non-social appetitive reward.

### Tonic firing of LC-NA is associated with distinct maternal behaviors

Pup retrieval is one of a suite of maternal behaviors exhibited by mothers and surrogate mice including nest building and maintenance, licking and grooming, crouching, etc. ^41^ Disruption of NA signaling in pregnant female rodents with neurotoxins, lesions, or genetic manipulation causes severe deficiencies in these behaviors postpartum, leading to pup negligence and elevated mortality ^24, 42, 43^. The relatively extended timeframe for our single unit recordings (40 – 90 min) afforded us the opportunity to monitor LC-NA firing patterns during a broader set of maternal behaviors beyond pup retrieval. By measuring tonic spiking as the mean log of instantaneous firing rate (IFR), we identified several maternal behaviors that were associated with sustained elevations in LC-NA activity relative to control conditions when the surrogate is alone in the arena (black dots in Figure 6A).

Nest building and maintenance are important components of maternal behavior in mice that become more intensive and elaborate in peripartum females ^44^. Mean IFR significantly increased while surrogates were organizing disturbed nesting material and covering pups with it (Fig. 6A, B). Importantly, mean IFR was not only higher than control periods, but it was also significantly higher than the IFR during episodes of nest maintenance when pups were not present in the nest and during digging, another active behavior (Fig. 6C, D). Licking and grooming pups is another fundamental maternal behavior that is critical for infant-mother bonding ^23, 45^. Although when surrogates licked pups, LC-NA single units did show a trend toward elevated FR, it never reached statistical significance as compared to the control period (prior to pup introduction to experimental arena, Fig. 6F). Importantly, this trend was not observed when surrogates gnawed on a toy mouse (Fig. 6G). Pup licking in Dbh-Cre surrogates injected with GCaMP6s, however, led to significant increase in calcium signal which persisted as long as the licking continued (Fig. H, I; n = 6 mice; licking: 0.32 ± 0.15 Z-score; control: −0.29 ± 0.04 Z-score; paired test, *p* < 0.05). This discrepancy could be due to the spatiotemporally integrative nature of calcium sensors, collecting fluorescence from throughout the neural population. These results are consistent with LC-NA influencing distinct maternal behavior patterns at short timescales and over longer intervals.

### Ca^2+^ signals in LC correlate with subsequent retrieval but not other locomotion

The onsets of phasic bursts in LC reliably preceded pup retrieval. To examine the temporal relationship between these two events in greater detail, we used automatic position tracking software (DeepLabCut) ^46^ to calculate a directionless instantaneous velocity signal (speed). Inspection of plots of the speed trace aligned to the concurrently recorded GCaMP signal revealed a peak in Ca^2+^ signals leading the peak in speed corresponding to the female’s return trip to the nest with the pup in its mouth by several hundred ms (Fig. 7A). No such Ca^2+^ transient was evident prior to other movements with similar temporal profiles.

**Figure 7:**
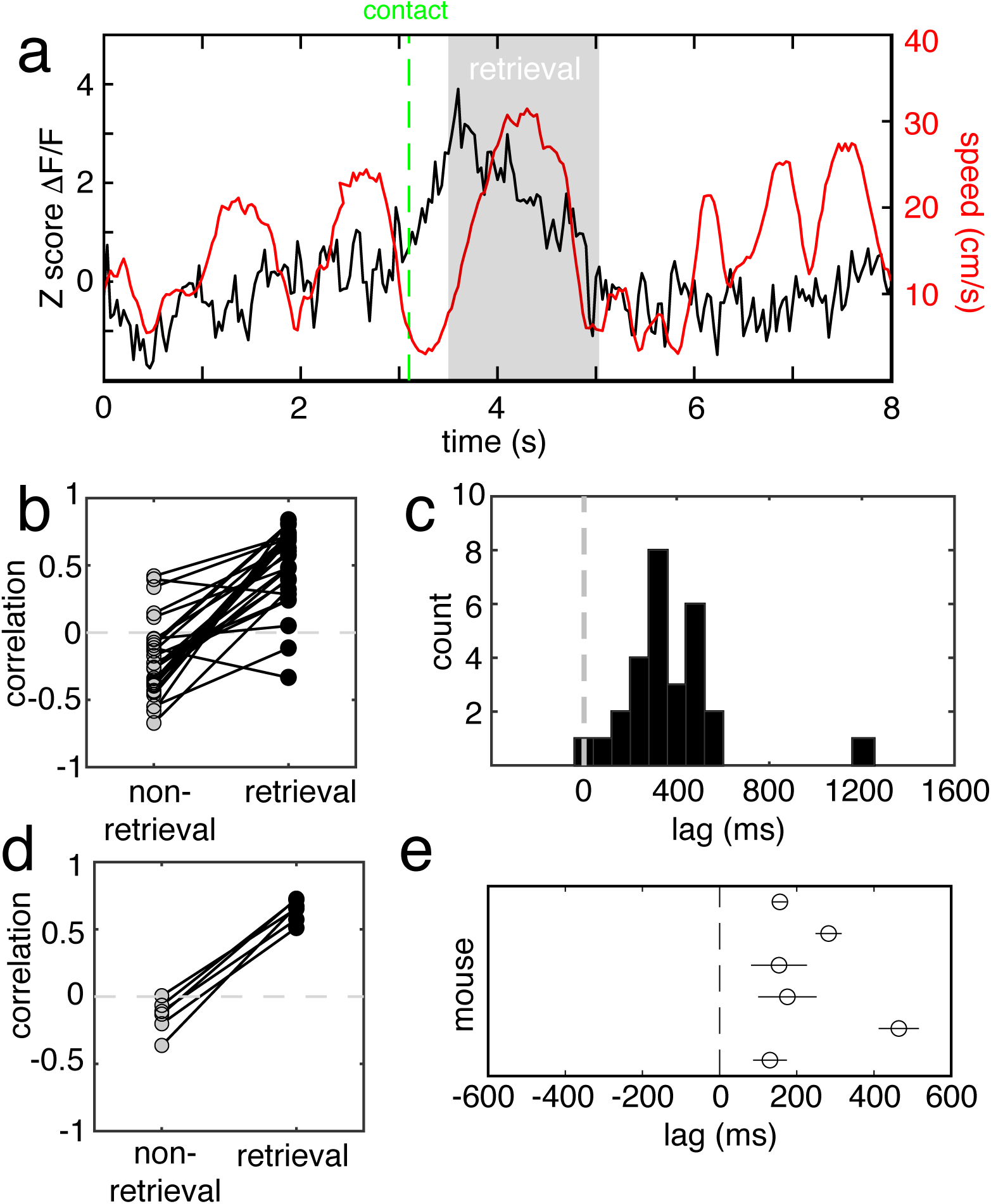
Ca^2+^ signals in LC correlate with subsequent retrieval but not other locomotion. (a): Plot of Z-score ΔF/F signal (black) and the concurrent speed signal computed from automated tracking (red) for a single retrieval trial. The dashed green line marks the point of contact with the pup, and the gray shaded region marks the time of retrieval, beginning at pup lift and ending in pup drop. This is a typical example showing that the fluorescence signal is unrelated to the animal’s speed at all times other than retrieval, including approaching the pup. (b): Plot comparing speed-ΔF/F correlation values between retrievals and pre-retrieval times for 28 different retrieval trials during one session from the mouse depicted in (a). Correlation values were significantly greater during retrieval than during non-retrieval (n = 28 trials; non-retrieval: −0.20 ± 0.05 retrieval: 0.73 ± 0.04; paired t test, p < 0.001) (c): Histogram of the latency between the ΔF/F signal and the speed signal for each trial in the session depicted in (a) and (b), computed as the max of the cross correlogram. The mean and SEM of the latency were 464 ± 52 ms. (d): Plot comparing the mean speed-ΔF/F correlation between retrievals and pre-retrieval times over all trials for 6 mice. The correlation values were significantly higher during retrieval (n = 6; baseline correlation: −0.14 ± 0.05; retrieval correlation: 0.61 ± 0.03; paired t test, *p* < 0.01). (e): Scatterplot of the mean ± SEM of the lag between the ΔF/F signal and the speed signal for each of the mice in (d).

We quantitatively verified this impression by performing for each retrieval trial a sliding cross correlation between the GCaMP signal and the mouse speed trace spanning a lag of −0.5 s to +*n* s for speed with respect to GCaMP (where the *n* is the duration of retrieval in seconds). We measured the lag and the Pearson correlation value at the peak of the cross correlation function, which we designated as the optimal lag. We compared this to the correlation values at the same lag for the prior 2.5 s of data. Figure 7B and C show the results of this analysis of all retrieval trials from the mouse depicted in Figure 7A. Comparison of the correlation values during retrieval to those before retrieval shows that mean correlation during retrieval was significantly higher (Fig. 7B; baseline correlation: −0.20 ± 0.05; retrieval correlation: 0.51 ± 0.06; paired t test, *p* < 0.001), and the histogram in Figure 7C shows that the optimal lag for all trials was short, consistent, and always GCaMP leading (lag = 460 ± 52 ms). This pattern was consistent across animals (Fig. 7D, E). Mean correlation values during retrieval were significantly higher than those at other times when the mouse was still making similar movements, including pup approach (n = 6; baseline correlation: −0.14 ± 0.05; retrieval correlation: 0.61 ± 0.03; paired t test, p < 0.01). Based on these data, we propose that LC’s role in action selection during maternal behavior is context-dependent and goal-directed.

## DISCUSSION

For decades, the LC-NA system has been understood to be a critical participant in arousal and attention in structured cognitive tasks. Specifically, LC neurons respond to novelty ^5^, behavioral significance ^3, 4^, effort ^39^, and changes in strategy ^6–8^, and they also promote goal-directed action selection ^9–13^. Largely independently, LC is also closely tied to social behaviors, such as individual recognition ^20–22^, mating choices ^18, 19^, and importantly for our work, maternal behavior ^23, 24^. Until now, it was unclear how these diverse functions are compartmentalized or multiplexed within the output of this very small population of neurons. It was also unknown at what time scale LC regulates natural behavioral interactions with conspecifics. Our results begin to provide the first answers to these questions, revealing an unexpectedly high level of precision in LC’s control of maternal behavior, and also highlighting some common principles that govern the role of phasic NA activity in both behavioral contexts.

Here we used chronic *in vivo* electrophysiology and fiber photometry to measure single unit firing and population activity in LC of female mice while they freely interacted with pups. Individual neurons in maternally experienced surrogates consistently showed a brief phasic burst near the initiation of pup retrievals. Optical recordings using genetically encoded calcium sensors restricted to expression only in LC-NA neurons confirmed this phasic response during pup retrievals, and further revealed it to be a pervasive event, reflecting relative synchrony among LC-NA. Phasic bursts in LC were not primarily driven by sensory responses to pups, because they were only seen on trials where the female actually lifted the pup, and they appeared at full magnitude on the first trials where the mouse began to retrieve. LC retrieval responses also did not simply reflect vigorous motor activity, general effort, or unexpected reward. Instead, they were most closely correlated with the specific, impending goal-directed action of returning the pup to the nest. At the same time, other sustained maternal behavioral states were reflected in the ongoing tonic LC activity (e.g. nest building in the presence of pups as opposed to the same behavior without pups). Therefore, we propose that LC regulation of maternal behavior shares a common blueprint with activity underlying structured tasks: Tonic firing levels encode slowly fluctuating state variables such as arousal and are punctuated by phasic events that promote goal-directed behavioral choices ^9–13^.

In our electrophysiology experiments, we observed two distinct types of cells that responded during pup retrieval. First, we observed putative LC-NA, that bore the characteristics of TH+/Dbh+ neurons and all increased firing during retrieval. Second, we saw another type of cell that responded with a sharp decrease in firing rate during retrieval. Because these cells were consistently anti correlated with LC-NA during retrieval, we speculate that they may represent some of the inhibitory neurons that are intermingled with LC or adjacent to it. Based on ultrastructural neuroanatomy, *in vitro* optogenetic activation, and *in vivo* manipulation of pupillary dilation, several groups have reported evidence of local inhibitory control of LC from different cell clusters. Since we can’t be sure about the precise location of the neurons we recorded, further work will be needed to clarify the specific involvement of any of these sources of inhibition.

Although LC was once viewed as a syncytium, firing largely in a highly coordinated fashion across the nucleus, recent evidence has cast doubt on this model. Closer examination of the anatomical distribution of inputs and outputs of LC suggests that there is greater specificity to them than previously appreciated, potentially forming separate parallel circuits that link specific afferent populations to specific targets ^15, 35^. Functional analysis of neuronal activity in multichannel recordings is consistent with this revised view, showing relatively little correlation among neurons ^16^. In contrast, our data seems to suggest that, at least under certain conditions, the population of LC-NA may act in close coordination, constituting a ‘broadcast’ signal that simultaneously mobilizes a wide swath of downstream circuits. This apparent synchrony is of course relative because individual LC-NA exhibited a diversity of activation latencies ranging from time of contact with the pup through the end of retrieval several seconds later. Indeed, the averaged firing from all LC-NA closely resembled the population response as measured by bulk Ca^2+^ signal. This phase diversity could arise from distinct sets of afferents driven by various behavioral components (sensory, motivational, motoric, etc). Further work dissecting distinct LC-NA ensembles by e.g. retrogradely labeling them from different efferent structures may clarify this. Nevertheless, we submit that the timescale of phasic events during maternal retrieval reveals a surprising level of precision and coordination among LC-NA.

We closely examined the temporal relationship between Ca^2+^ signals and locomotion during retrieval. We found that LC activity consistently preceded the change in speed associated with initiating retrieval by several hundred ms. LC fluorescence was highly correlated with speed during retrieval periods, yet these two measures were either not or negatively correlated during pup approach. Several recent studies reported significant correlations between phasic activation of LC and motion on approach to a learned reward location ^47, 48^. One interpretation of our results that is concordant with these previous findings is that an important feature of LC’s regulation of maternal behavior is the promotion of actions that achieve the motivational goal of delivering the pup to a specific reward-associated location (the nest).

In conclusion, we propose that our results begin to integrate and align the participation of LC-NA in both unstructured natural social interactions and experimenter-designed and instructed cognitive tasks into a common framework. Moving forward, we are optimistic that our results delineate a novel approach for deconstructing the ethological building blocks that have been adapted to solve closely controlled problems devised for the laboratory setting.

## METHODS

### Animals

All procedures were conducted in accordance with the National Institutes of Health’s Guide for the Care and Use of Laboratory Animals and approved by the Cold Spring Harbor Laboratory Institutional Animal Care and Use Committee. Adult (8-12 weeks) female mice were used in all experiments. C57Bl/6 mice (The Jackson Laboratory) were used for single unit electrophysiology experiments and Dbh-Cre mice (Tg(Dbh-cre)KH212Gsat/Mmucd, unfrozen stock, MMRRC) were used for fiber photometry experiments. Animals were maintained on a reversed 12h/12h light/dark cycle (lights off 09:00), and All experiments were performed during the animals’ dark cycle. Food and water were available *ad libitum*.

### Genotyping

Hemizygous male Dbh-Cre were crossed with Wild Type (C57Bl/6) females. The resulting offspring were genotyped according to the vendor protocol (MMRRC). After weaning on postnatal day 21, tail samples were collected under brief isoflurane anesthesia. Samples were dissolved in lysis buffer (10 mM NaOH and 0.1 mM EDTA) and proteinase K at 37°C for 4 h, and the proteinase K was deactivated in a 95°C water bath. The PCR solution included 1 µl of the DNA solution, 10µl PCR master mix (Promega; GoTaq Green Master Mix M7123), 7µl nuclease free water, and 1µl of each primer (10µm, 5’: FAATGGCAGAGTGGGGTTGGG, 3’: CGGCAAACGGACAGAAGCATT, Sigma-Aldrich, USA).

### Viruses

Cre-dependent adeno-associated virus (AAV; serotypes 5 or 9) was used to express GCaMP6s (AAV5-Syn-Flex-GCaMP6s, Addgene), or GCaMP7f (AAV9-Syn-Flex-jGCaMP7f-WPRE, Addgene) in the LC of Dbh-Cre mice ^49, 50^.

### Surgery

All surgeries were performed under isoflurane anesthesia (2%-3% induction, 0.7%-1.2% maintenance) in a stereotaxic frame. To reduce pain and inflammation, mice were injected with meloxicam (5mg/kg) prior to all surgeries and meloxicam gel (ClearH2O) was added to singly housed mice after the surgery.

For electrophysiology implants, a ∼1 mm x 1mm craniotomy was made above the estimated LC location (from Bregma, AP: −5.1 mm – −5.4 mm, ML: 0.8 mm – 1.0 mm). LC was first located using a single tungsten electrode (1MΩ, Microprobes) by well-established neurophysiological criteria (slow tonic firing, wide spike shape, phasic response to tail pinch) (Shea et al, 2008). After successful location of LC, an 8 or 16-channel movable microwire bundle was attached to 18 pin Omnetics connector (Innovative Electrophysiology) and was implanted with the tip of the bundle ∼800µm above LC. The implant was secured to the skull with adhesive luting cement (Parkell, Inc.). For additional support, two machine screws (Amazon Supply) were secured to the skull and the ground wire was looped around both. Additional luting cement was then applied to cover and secure the implant. Mice were allowed to recover for 7 day before the bundle was advanced.

For fiber photometry implants, a similar craniotomy was made at the same position. Injection of a Cre-dependent AAV driving expression of the Ca^2+^ sensor GCaMP (either AAV5-Syn-Flex-GCaMP6s or AAV9-Syn-Flex-jGCaMP7f-WPRE) was slowly (∼100 nl/min) injected local to LC (from Bregma, AP: −5.2mm, ML: 0.85mm, DV: 2.9mm from brain surface). After completion of the injection, the injection pipette was left in place for an additional 5min prior to withdrawal. Subsequently, a 200 µm diameter optic fiber (NA 0.37, Doric Lenses) was implanted slightly above the injection site and LC (DV: 2.7mm from brain surface) and was cemented in place. Mice were allowed to recover and express GCaMP for at least three weeks before experiments began.

### Behavior

In most cases, surrogates were nulliparous female mice that were co-housed with primiparous CBA/CaJ females (The Jackson labs) beginning 1 – 5 days before delivery. A subset of surrogates were exposed to the same litter of pups for only 30min a day through PND 5 days (Fig. 4). The same subset was injected with GCaMP7f. All behavior was conducted in the dark, in a Plexiglass arena (42 cm x 28 cm) inside a custom-built double-walled anechoic chamber (IAC). To reduce anxiety, mice were introduced to handling and experimental arena for at least 1 h/d for a week prior to commencing experiments. Bedding in the arena was unchanged through the end of the experiment. Pup retrieval behavior was elicited as follows. Each electrophysiology recording session had a duration of 40 – 90min. The sessions began with the surrogate alone in the arena. After a short habituation period, the entire litter of pups (5 – 10) were scattered around the arena and allowed to be retrieved back to nest by the surrogate. This procedure was repeated several times during each session (approx. every 10 minutes). Between each retrieval surrogates were allowed to freely interact with the pups. Fiber photometry sessions were similar, but shorter (∼10 min), and pups were only scattered once. Post hoc scoring of various behaviors was conducted manually using BORIS ^51^ from recorded videos (30fps). Pup contact was defined as the first frame in which mouse’s snout was a top of a pup. Retrieval onset was defined as the first frame in which the pup was lifted from the bedding.

### Histology and Immunohistochemistry

Mice were perfused with 4% paraformaldehyde/PBS, and brains were extracted and post-fixed overnight at 4°C. Brains were then treated with 30% sucrose/PBS overnight at room temperature (RT) and sectioned on a freezing microtome at 50µm. For fiber photometry subjects, free-floating sections were immunostained using standard protocols. Briefly, sections were blocked in 5% normal goat serum + 2% BSA and 2% Triton-X for 1 h and incubated with the following primary antibodies overnight at 4°C: rabbit anti-GFP (1:1000, Invitrogen) and chicken anti-TH (1:1000, Alves Labs). The next day, the sections were washed in PBS and incubated with secondary antibodies (Alexafluor 488nm goat anti-rabbit, 1:500 and Alexafluor 594nm goat anti-chicken, 1:500; Invitrogen) for an additional 1 h and mounted in Fluoromount-G (Southern Biotech). For mice implanted with wire bundles, sections were Nissl stained with cresyl violet to identify the deepest location of the bundle. Images were acquired using an Olympus BX43 microscope (X4 or X10 objective, UPlanFL N). Mice that either had a misplaced optical fiber or wire bundle or inadequate viral expression were excluded from the study.

### *In vivo* electrophysiology

The electrode bundle was advanced daily by 50 – 100 µm until putative LC neurons were reached (defined by depth and known electrophysiological properties as described previously, including Shea et al., (2008)). Once the bundle reached the location of LC, co-housing of the implanted mouse with a pregnant CBA/CaJ female began, and the surrogate was introduced to daily behavioral tests as described above. Electrodes were connected by a tether to a headstage and amplified (Tucker-Davis Technologies). This allowed the mice to move freely while performing natural behaviors. Recordings were digitized at 24 kHz, bandpass-filtered from 300Hz to 3kHz and thresholded online by the user and saved for further analysis using Synapse software (Tucker-Davis Technologies). Posthoc spike sorting was first automatically performed using OpenSorter software (Tucker-Davis Technologies) and further refined manually. Standard criteria were used to ensure single unit isolation (e.g manual inspection of cluster separation, SNR, and autocorrelation histograms).

### Fiber photometry

GCaMP signals were detected and measured as follows. A 200 µm optical fiber cable (NA 0.39) was mated to the fiber implant at the beginning of each optical recording session, and it was used to deliver 470 nm and 565 nm excitation light to the brain. The intensity of the light for excitation was adjusted to ∼30 µW at the tip of the patch cord. The two wavelengths were sinusoidally modulated 180 degrees out of phase at 211 Hz. Green and red emitted light signals were filtered and split to separate photodetectors and digitally sampled at 6100 Hz via a data acquisition board (National Instruments, model #NI USB-6211). Peaks were extracted by custom Matlab (Natick, MA) software to achieve and effective sampling rate of 211 Hz. Each signal was separately corrected for photobleaching by fitting the decay with a double exponential. A robust regression algorithm was used to compute the coefficients of a linear transformation between the red and green signal, which was then applied to the red signal to generate a prediction of the green trace (G_p_). This predicted trace was subtracted from the measured green trace and the residual (G_r_) was used to calculate ΔF/F according to the following equation:

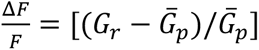

The resulting traces from each recording session were collectively converted to a z-score to compare between subjects.

### Data analysis

Unless specified otherwise all data analyses were performed in Matlab (MathWorks, USA) using custom written code. Recordings from single units were sorted into three groups according to defined electrophysiological characteristics: (1) Putative LC units exhibited a wide spike shape (1.5 – 2ms), slow (0.5 – 5 Hz) regular tonic firing rates, no jaw related firing events (e.g while gnawing on corncob bedding), and phasic responses to salient stimuli (e.g. a sudden sound, the experimenter’s hand, etc.); (2) Inhibited units exhibited narrow spike shapes (0.7 – 0.9 ms), fast (>5 Hz) irregular (CV = 2.07 ± 0.76) tonic firing rates, and strong firing rate suppression during pup retrieval; (3) Unidentified units exhibited heterogenous characteristics and could not be clearly separated into one of the other groups. We cannot rule out the possibility that some unidentified cells are LC neurons. Peri-stimulus time histograms (PSTHs) were generated for 20 s periods (±10 s from retrieval onset) by binning spike rates into 0.2 s bins. Individual cell PSTHs were z-scored and heatmaps of averaged z-score were calculated. Max z-scores for baseline (−2.8 s – −0.8 s from behavior onset) and behavior (pup retrieval, toy mouse retrieval, snack; −0.8 s – 1.6 s from behavior onset) were extracted.

Instantaneous firing rate was calculated as the inverse of each inter-spike interval (ISI). To perform correlations between simultaneously recorded neurons, we binned the spike train of each neuron into 1 s bins. The correlation value at each time point was computed as a Pearson correlation between the bins of each neuron for a 30 s window centered around that time point. Retrieval trials were shuffled by randomly permuting trial identities for one of the pair of neurons. Non-retrieval activity was shuffled by taking the mean correlation value for all time steps as one set of bins was circularly permuted in time with respect to the other.

DeepLabCut ^46^ was used to track the position of the mouse throughout each recording session. We achieved the most reliable tracking by training DLC to follow the connection of the tip of the patch cord to the fiber. The XY coordinates returned by automated tracking were used to compute a directionless velocity signal (speed), which was then smoothed by convolution with a 7-point boxcar kernel. ΔF/F were resampled at 30 Hz to match the video frame rate. Optimal lag for speed relative to ΔF/F was found by calculating the maximum of the cross correlogram of speed versus the ΔF/F signal from 0.5 s prior to retrieval onset through the end of retrieval (when the pup was dropped in the nest). Speed was shifted accordingly and Pearson correlation were calculated for the baseline period (−2.5 s – −0.5 s prior to retrieval) and retrieval (−0.5 s – retrieval offset).

## Acknowledgements

The authors wish to thank C. Kelahan, J. Sturgill, and L. Huang for technical assistance, and Shea Lab members for helpful comments and discussion. This work was supported by a grant to SDS from the National Institute of Mental Health (R01MH119250) and a National Alliance for Research on Schizophrenia and Depression Young Investigator Grant to RD from the Brain and Behavior Research Foundation

## Author Contributions

SDS supervised the project and designed the experiments together with RD. RD developed the methods and collected the data. SDS and RD analyzed the data and wrote the paper.

---

**Extended Data Fig. 1:**
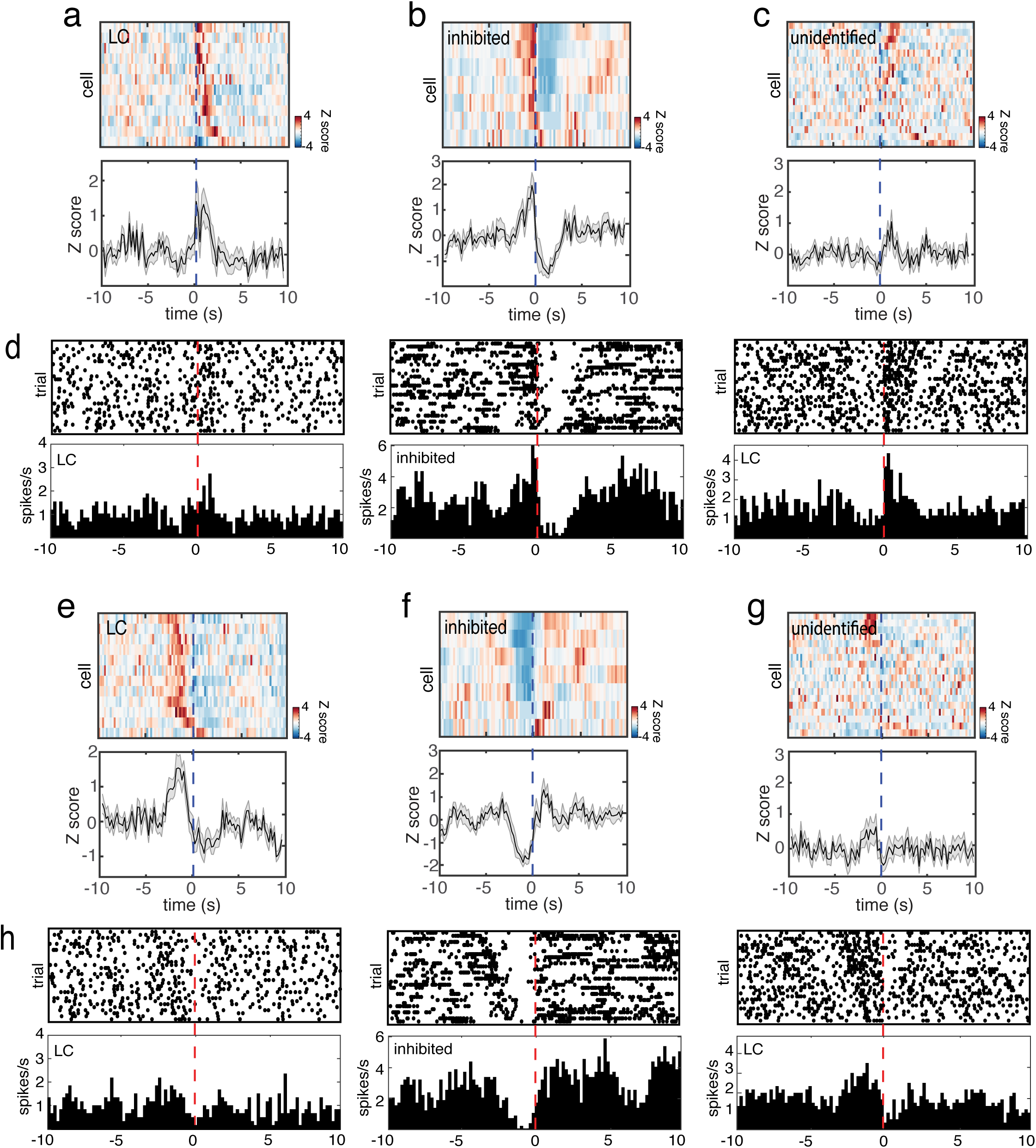
Retrieval Inhibited (RI) neurons show an increase in firing rate prior to pup contact and strong activity rebound following pup drop. Same data as in Fig. 2a-f aligned to pup contact (a-d) or pup drop (e-h).

**Extended Data Fig. 2:**
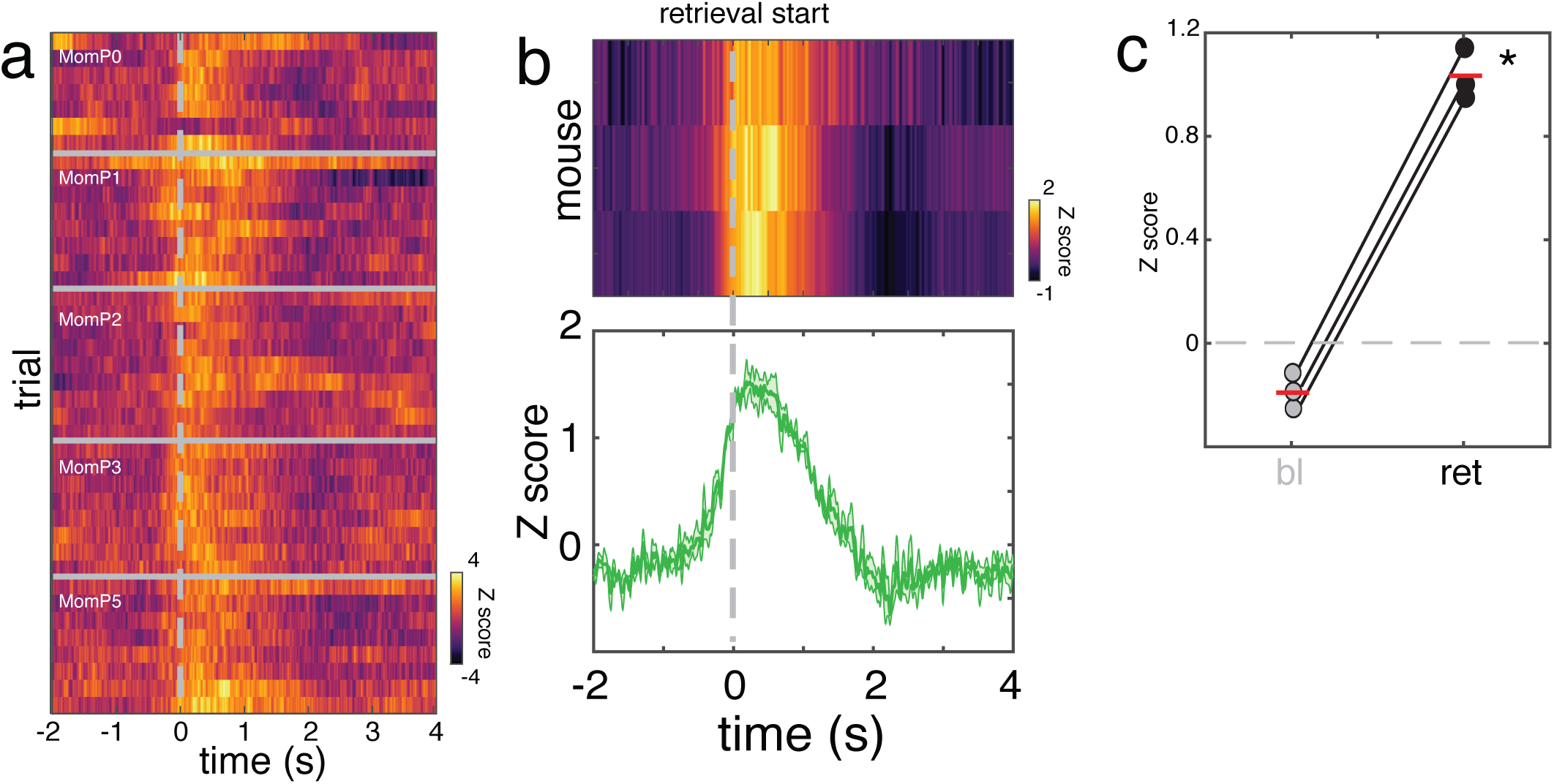
Fiber photometry in LC of mothers during pup retrieval shows pervasive burst similar to surrogates. (a): Example of bulk Ca2+ signal fluctuations measured during pup retrieval in the same mother over 6 days (n=40 individual trials; PND0 – PND5) aligned to pup lift (dashed line). The same female was previously used as surrogate (Fig. 3e). The lower SNR is probably due to degradation of the signal after close to 3-month post injection. (b): Mean Z-scored ΔF/F responses across all retrievals aligned to pup lift (n=3 mothers; 1 GCaMP6s, 2 GCaMP7f; *top panel*) and the mean ± SEM responses across all mothers. (c): Scatterplot comparing the baseline activity (bl, grey circles) to the retrieval activity (ret, black circles). Red lines represented the mean of each condition (baseline: −0.18± 0.07; retrieval: 1.03± 0.1; paired t-test; *p* < 0.001).

